# Multitasking practice eliminates modality-based interference by separating task representations in sensory brain regions

**DOI:** 10.1101/2024.03.26.586741

**Authors:** Marie Mueckstein, Kai Görgen, Stephan Heinzel, Urs Granacher, Michael A. Rapp, Christine Stelzel

## Abstract

The debate on the neural basis of multitasking costs evolves around neural overlap between concurrently performed tasks. Recent evidence suggests that training-related reductions in representational overlap in fronto-parietal brain regions predict multitasking improvements. Cognitive theories assume that overlap of task representations may lead to unintended information exchange between tasks (i.e., crosstalk). Modality-based crosstalk was suggested as a source for multitasking costs in multisensory settings. Robust findings of increased costs for certain modality mappings may be explained by crosstalk between the stimulus modality in one task and sensory action consequences in the concurrently performed task. Whether modality-based crosstalk emerges from representational overlap in general fronto-parietal multitasking regions or modality-specific regions is not known yet. In this functional neuroimaging study, we investigate neural overlap in multitasking performance, focusing on modality compatibility by employing multivariate pattern analysis and modality-specific practice interventions in three groups (total N = 54). We observed significant differences between modality compatible and modality incompatible single-task representations, specifically in the auditory cortex but not in fronto-parietal regions. Notably, improved auditory decoding accuracy related to modality incompatible tasks was predictive of performance gains in the corresponding dual task along with complete elimination of modality-specific dual-task costs. This predictive relationship was evident only in the group practicing modality incompatible mappings, suggesting that specific practice on task sets with modality overlap influenced both neural representations and subsequent multitasking performance. This study contributes to the integration of cognitive theory and neuroscience and the role of task representations in dual-task interference.

**Significance:** In a society dominated by multitasking, understanding its neurocognitive basis and plasticity is crucial for key aspects of work-related and everyday tasks. We investigate the neural mechanisms behind multitasking limitations, offering insights for targeted cognitive interventions. The study builds upon established theories of cognitive multitasking and imaging research, addressing the concept of modality-based crosstalk - the unintended exchange of modality-based information between tasks. Through functional brain imaging and pattern analysis, we examined how neural task representations contribute to performance costs in dual tasks with varying degrees of modality overlap. Notably, our findings demonstrate a practice-related decrease in neural overlap which is associated with substantial multitasking improvements, specifically in the auditory cortex, emphasizing the contribution of sensory regions to flexible multidimensional task representations.

## Introduction

Human limitations in multitasking are significant and can lead to safety-relevant consequences in everyday life, for example, when using a mobile phone while driving. A long-held debate relates to the question of whether performance costs in multitasking emerge based on the neural overlap of concurrently performed tasks (Just et al., 2001; Klingberg, 1998). Recent multivariate imaging research supported this idea, revealing that multitasking training reduces the overlap of concurrent task representations in fronto-parietal brain regions which predicts training improvements in multitasking (Garner & Dux, 2015). According to cognitive multitasking theories, the overlap of task representations can result in the non-intentional exchange of information between tasks, also called central crosstalk (Janczyk et al., 2014; Koch, 2009; Logan & Gordon, 2001; Navon & Miller, 1987). Crosstalk may lead to between-task benefits or interference at different levels of a task, which is supported by behavioral and neural research (Halvorson & Hazeltine, 2015; Koch et al., 2018; Lien & Proctor, 2002; Paas Oliveros et al., 2023).

Recently, modality-based crosstalk was suggested to underlie increased multitasking costs when comparing dual tasks with different modality mappings (Hazeltine et al., 2006; Schacherer & Hazeltine, 2020). For example, visual-manual and auditory-vocal (i.e. modality compatible) modality mappings produce consistently lower dual-task costs than visual-vocal and auditory-manual (i.e. modality incompatible) mappings (Figure 1B) (Göthe et al., 2016; Hazeltine et al., 2006; Stelzel et al., 2006). Modality-based crosstalk refers to the interference between the stimulus modality in one task and sensory action consequences (i.e. action effects) in the concurrently performed task, while the presented stimulus and required response modalities do not overlap between tasks.

**Figure 1.**
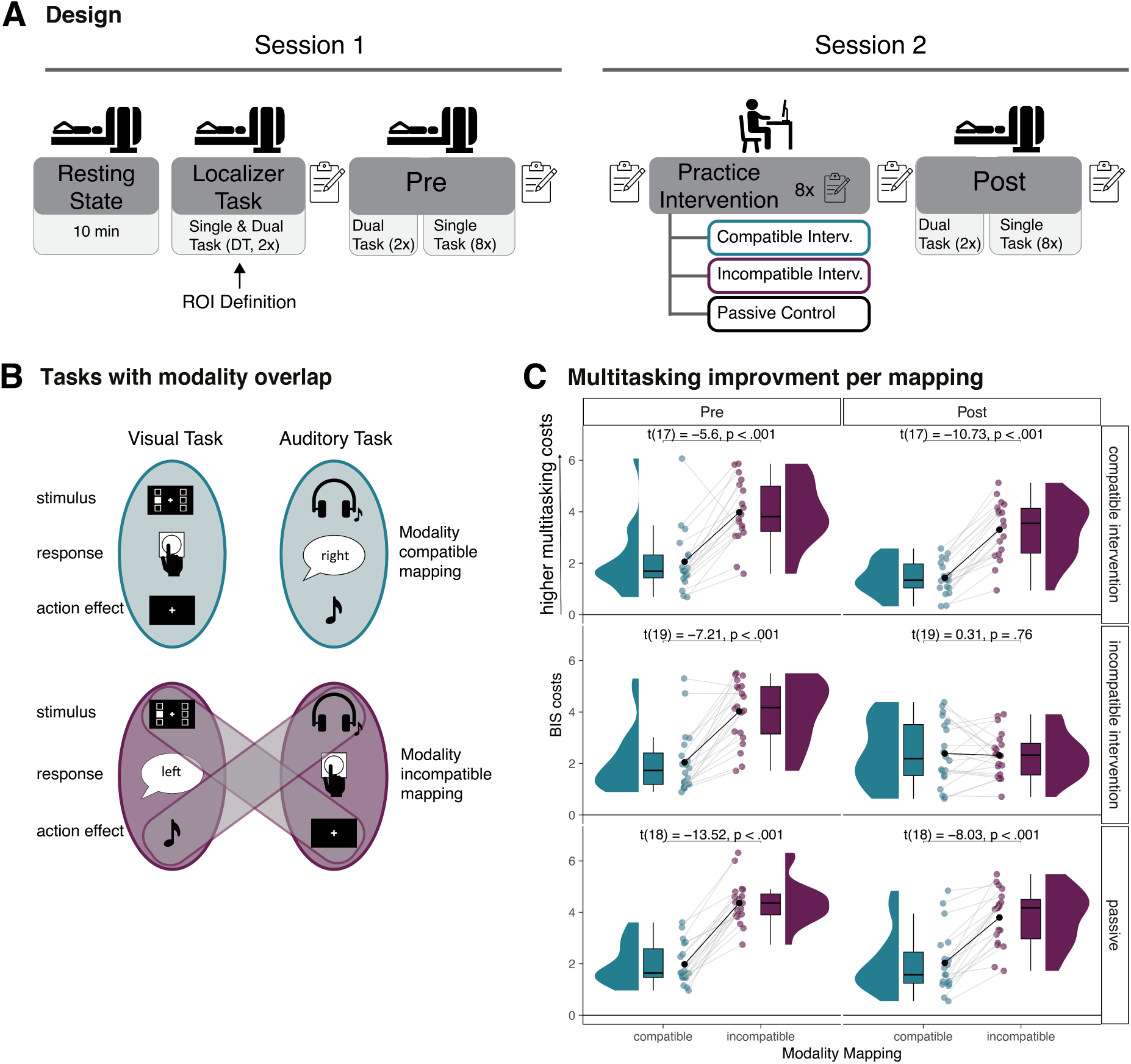
Study design, modality mappings and behavioral results. *Note:* **A**: Overall study desgin with two sessions. Resting state data and subjective items (before and after each part) will be reported elsewhere. **B**: The upper part of the figure shows the stimulus-response pairings for the modality compatible mapping, comprising of a visual stimulus with the manual response and the auditory task combined with the vocal response. For each response, the corresponding action effect is depicted as well. Note that the action effect of the manual response is not exclusively visual but also somatosensory. The lower part depicts the modality incompatible mapping. The visual stimulus is paired with a vocal response and the auditory stimulus with a manual response. In this condition, the match between action-effect modality and stimulus modality is between tasks, potentially causing interference due to higher overlap. **C**: The graph shows distributions, boxplot, mean (black dot) and individual performance per modality mapping, timepoint and intervention group. Almost all individuals show the robust difference of multitasking costs between modality compatible and modality incompatible mapping. Multitasking cost is the difference between dual task and single tasks, measured as BIS, which is an integration of reaction times and accuracies, BIS = 0 means no difference between single and dual task performance). Comparing the Pre and Post timepoint, only the incompatible intervention group eliminated this robust difference after the practice intervention (middle panel on the right). All pairwise t-tests are corrected with the Benjamini-Hochberg procedure (Benjamini & Hochberg, 1995).

So far, the modality-based crosstalk assumption has been supported primarily by behavioral research (Göthe et al., 2016; Schacherer & Hazeltine, 2020, 2023). It remains unclear how modality-based crosstalk evolves at the neural level and how this is affected by multitasking practice. Specifically, it is still to be determined if the assumed overlap of stimulus modality and action-effect modality is also present on a neural level and if so, whether representational overlap is present in general multitasking-related brain regions (i.e., fronto-parietal regions, as identified by previous research on multitasking training (Garner & Dux, 2015) and by a meta-analysis on the neural locus of multitasking (Worringer et al., 2019)) or rather in modality-specific sensory brain regions (i.e., visual or auditory regions). Consequently, here we ask how the potential overlap of neural representations contributes to performance costs in multimodal multitasking settings, using dual tasks with different degrees of modality overlap between tasks. If such an association exists, we expect modality-specific dual-task practice to reduce this overlap. By comparing two dual tasks with identical stimuli and responses, but a different degree of modality overlap regarding sensory action effects, we are able to investigate whether and where in the brain the assumed overlap of task representations differs between modality mappings and whether it affects task performance.

In this functional Magnetic Resonance Imaging (fMRI) study, we used multivariate pattern analysis (MVPA) to decode single-task-related representations. We compared the effect of task practice on decoding accuracies within and between tasks, with and without modality overlap regarding action effects. The task combination of visual-manual and auditory-vocal (i.e., modality compatible mapping) is considered to have no modality overlap between stimuli and action effects. The visual-vocal and auditory-manual task combination (i.e., modality incompatible mapping) exhibits modality overlap (Figure 1B). We hypothesized that neural overlap is higher between tasks with a modality incompatible mapping in sensory brain regions related to the sensory action effects of the responses. Additionally, we expect this modality overlap to essentially contribute to multitasking performance, even though the concurrently performed tasks have different sensory inputs (i.e., visual and auditory). As the action effects of vocal responses are auditory in nature, we expected the most pronounced effects in auditory brain regions. Action effects for manual responses, in contrast, are multimodal, including visual and somatosensory modalities (Straube et al., 2017). Further, based on the training literature (Garner & Dux, 2015), we predicted a practice-related decrease of neural overlap in sensory brain regions, specifically for modality incompatible dual tasks. Participants completed single and dual tasks during fMRI measurements before and after a dual-task practice intervention (Figure 1A). The sample was randomly split into three different intervention groups (modality compatible intervention, modality incompatible intervention, passive control), two of them completing the same forced-choice dual tasks as during fMRI measurements for 80 minutes with visual and auditory input and manual and vocal responses, but varying in the amount of modality overlap with action effects between tasks.

Previewing our results, we replicate the complete elimination of the substantial difference in behavioral dual-task costs between the modality mappings after practicing the modality incompatible mapping (Mueckstein et al., 2022). Using MVPA on blood-oxygen-level-dependent (BOLD) response patterns in the single tasks, we found a significant difference in decoding accuracy between the modality incompatible mapping compared to the modality compatible mapping in the auditory region of interest, which was defined by univariate task activation in a localizer task (Figure 1A). This difference supports the assumption that the modality-related sensory task representations of the component tasks overlap more in the modality incompatible mapping compared to the modality compatible mapping, thus extending previous behavioral and neural findings in the field. Additionally, only for participants who completed the modality incompatible practice, a selective decrease in the neural overlap in the auditory cortex between the modality incompatible tasks was positively associated with substantial performance improvements.

## Results

### Influence of Modality Overlap on Behavior and Practice-related Changes

We assessed behavioral performance as a balanced integration score (BIS) of reaction times and accuracies (Liesefeld & Janczyk, 2019) (see Table 1 for reaction times and Table 2 for error rates), to account for the dependence of the two parameters and to have a single parameter for the following correlational analyses with the neural decoding parameter. Dual-task costs were calculated as the difference between single and dual tasks, in which higher values of the BIS parameter indicate higher dual-task costs (i.e., higher reaction times and error rates in dual tasks than in single tasks).

**Table 1.**
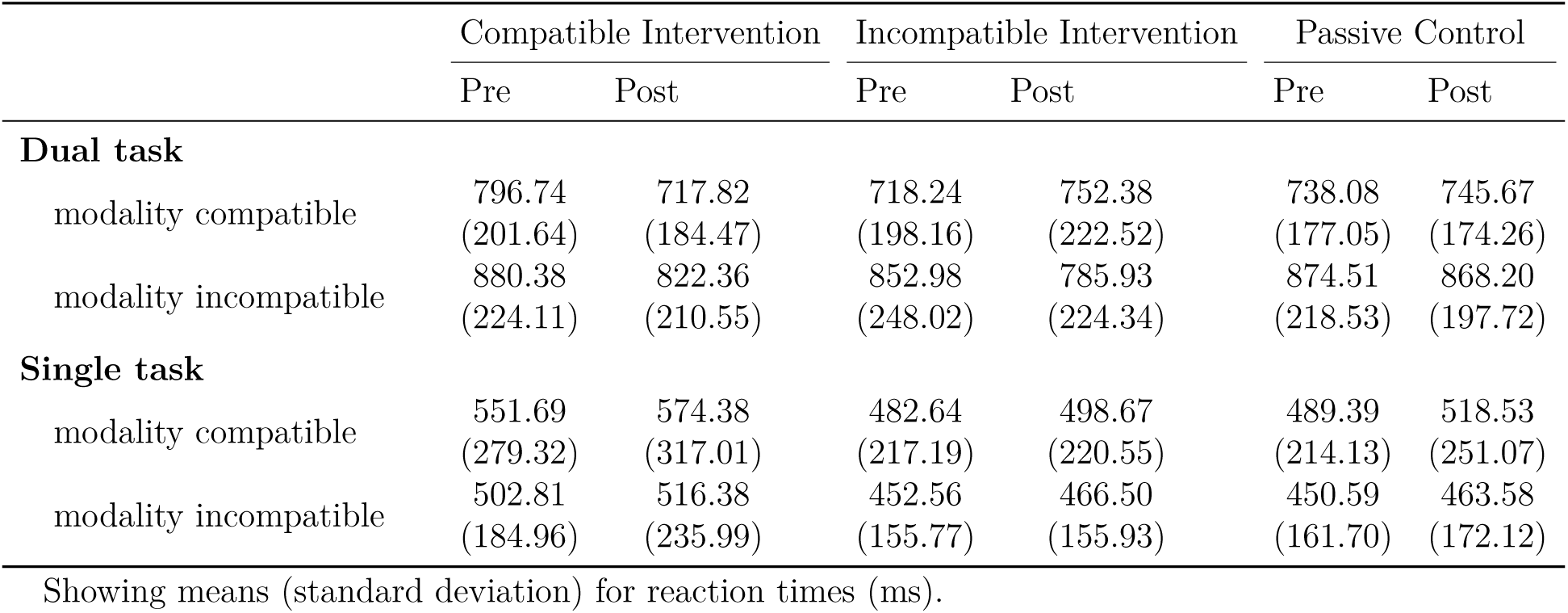
Raw reaction times per task type, modality mapping, timepoint and intervention group.

**Table 2.** Error rate per task type, modality mapping, timepoint and intervention group.

|  | Compatible Intervention |  | Incompatible Intervention |  | Passive Control |  |
| --- | --- | --- | --- | --- | --- | --- |
|  | Pre | Post | Pre | Post | Pre | Post |
| **Dual task** |  |  |  |  |  |  |
| modality compatible | 15.97<br>(13.59) | 15.49<br>(11.57) | 15.82<br>(13.00) | 23.55<br>(18.10) | 14.14<br>(6.81) | 17.84<br>(20.05) |
| modality incompatible | 37.59<br>(17.77) | 34.03<br>(18.19) | 37.15<br>(16.53) | 16.45<br>(10.56) | 41.94<br>(16.46) | 36.47<br>(23.24) |
| **Single task** |  |  |  |  |  |  |
| modality compatible | 1.99<br>(2.70) | 3.56<br>(6.95) | 1.37<br>(2.73) | 4.54<br>(10.21) | 1.36<br>(2.18) | 3.23<br>(7.68) |
| modality incompatible | 1.16<br>(2.00) | 3.77<br>(6.83) | 1.25<br>(2.46) | 3.63<br>(7.33) | 0.85<br>(1.57) | 3.64<br>(9.70) |

Showing means (standard deviation) for error rates in %.

At baseline, all three groups showed a robust behavioral effect of modality mapping, *F* (1, 54) = 178.92, *p < .*001, 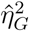 = .483, 90% CI [.323*, .*603] with higher dual-task costs for the modality incompatible mapping (*M* = 4.12, *SE* = 0.15) compared to the modality compatible mapping (*M* = 2.03, *SE* = 0.14). The difference between modality mappings was selectively eliminated only after the modality incompatible practice intervention. Omnibus Anova showed a significant interaction between the factors intervention group, time point and modality mapping *F* (2, 54) = 10.49, *p < .*001, 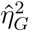 = .039, 90% CI [.000*, .*133]. Post-hoc tests confirmed that the difference between modality compatible and modality incompatible mapping was significant for all groups and time points, except for the incompatible intervention group after practice (see Figure 1C). This replicates our previous findings (Mueckstein et al., 2022) and indicates that modality-based crosstalk can be eliminated with specific practice.

### Neural Overlap of Task Representations

We investigated the overlap of task representations at baseline in task-relevant regions with MVPA on subject-specific beta images from first-level single task analysis. We trained linear support vector machines (SVMs) using a leave-one-run-out cross-classification (implemented in The decoding toolbox (Hebart et al., 2014)) to distinguish between the two single tasks from fMRI activity pattern in the modality compatible and in the modality incompatible mapping, respectively (compare Figure 1B). Task-relevant regions were defined by task-related univariate clusters in two separate localizer runs in which participants performed the same single and dual tasks as in the main experiment in a block design (Figure 1A. We contrasted the input modalities (visual vs. auditory), the output modalities (manual vs. vocal), and single vs. dual tasks (frontal region as multitasking specific region) to create five task-relevant clusters per hemispheres (grey cluster in Figure 2A) as a basis for the ROI analysis. Within each cluster, we defined a sphere (10 mm radius) centered at the maximum voxel on the group level (orange sphere in Figure 2A and peak coordinates in Table 5). Additionally, we added the fronto-parietal-subcortical cubic ROIs defined in a previous study on multitasking training (Garner & Dux, 2015) to compare our task-specific ROIs with those task-independent ones.

**Figure 2.**
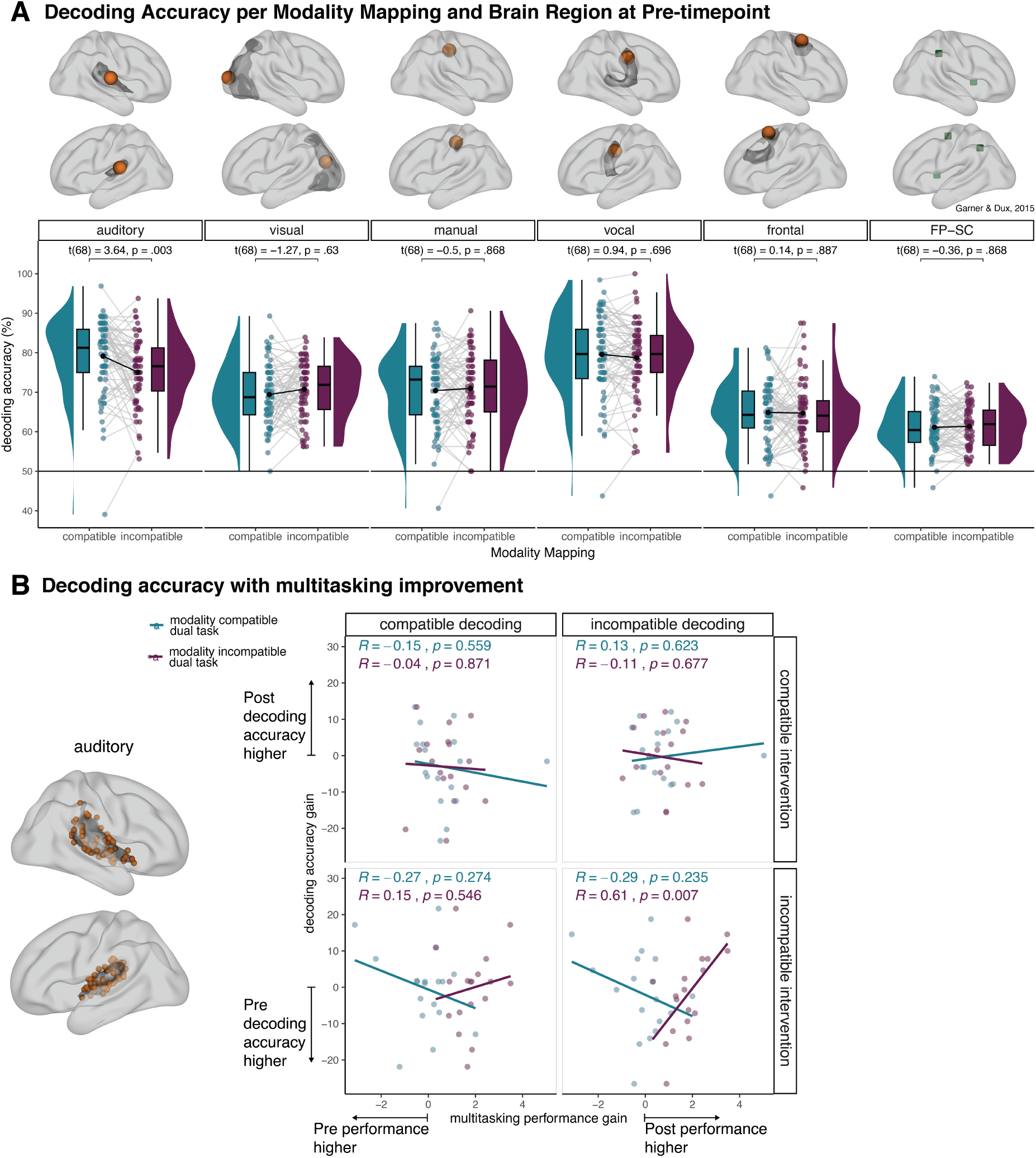
Neural task representation and the relation to performance. *Note:* **A**: Decoding accuracy per modality mapping and brain region at Pre timepoint. Group-based clusters are shown in grey and corresponding group spheres (10mm radius) in orange. FP-SC ROIs are defined by Garner & Dux (2015) as cubic ROIs. The graphs show distribution, boxplot, mean (black dot and line) and individual values. The decoding accuracy in the auditory ROI for the decoding between visual-manual and auditory-vocal tasks (modality compatible) are significantly higher compared to the decoding between visual-vocal and auditory-manual tasks (modality incompatible). All pairwise t-tests are corrected with the Benjamini-Hochberg procedure (Benjamini & Hochberg, 1995). Similar effects were found on cluster and whole-brain level (see Figure 3). **B**: Correlation between multitasking improvements and decoding accuracy gains in the auditory region. Orange spheres (radius 10mm) indicate individual peak activation within the group-based cluster. The correlation was only significant for participants who completed the modality incompatible intervention. Their multitasking performance improved from Pre to Post if they showed a reduced overlap between incompatible mapping tasks, indicated by higher decoding accuracy at the Post timepoint compared to Pre.

Our results demonstrate that the trained classifiers can robustly distinguish between the two single tasks (i.e., visual-manual vs. auditory-vocal and visual-vocal vs auditory-manual, respectively) in both modality mappings in all the task-relevant regions, decoding accuracies for all ROIs are significantly above chance level (50%), all corrected t-tests *p < .*001. Remarkably, a pairwise t-test, corrected for multiple comparisons, revealed a significant difference between the representation of the modality mappings only in the auditory region, *t*(68) = 3.64, *p* = .003. Here, decoding accuracies were higher for modality compatible tasks (*M* = 79.13%, *SE* = 1.16) compared to modality incompatible tasks (*M* = 75.07%, *SE* = 1.02). Note that the tasks only differed in the specific modality mappings - sensory inputs for the two mappings were identical, thus excluding pure bottom-up effects.

To rule out that the results are only due to our ROI selection, we ran the same analysis in the whole group cluster (Figure 3A), using a searchlight analysis on the whole brain (Figure 4) and tested for potential influences of task difficulty (Figure 3B) and task instruction (Figure 3C). All analyses confirmed the difference in the auditory region, except on the cluster level where the difference is only numerical (*p* = .34). Accordingly, the results in the following results sections will focus on this auditory region, which selectively differentiates between the modality compatible and modality incompatible mappings. We further investigated whether the higher neural overlap at baseline is also associated with behavioral performance. Surprisingly, we found no significant correlation between the decoding accuracy and the dual-task costs at baseline, *r*[-0.15, 0.03], *p >* 0.27. This could be due to two points: Firstly, there might be low reliability of estimates for the behavioral performance in the Pre session. Secondly, the strength of decoding alone might have no primary relevance for the behavior.

**Figure 3.**
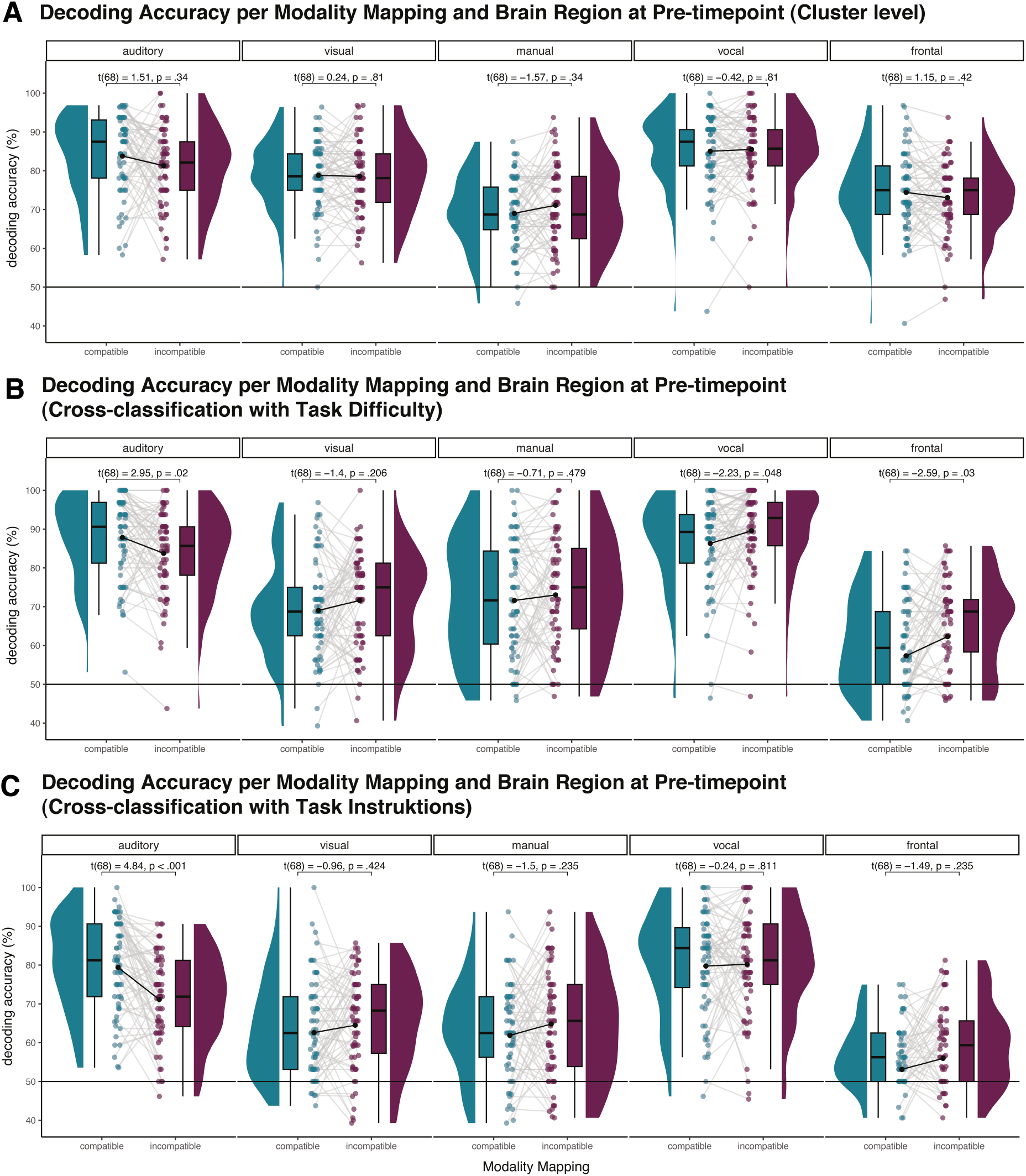
Control analysis for pattern analysis. *Note:* **A**: Decoding accuracy per modality mapping and brain region at Pre timepoint. Decoded on group-based cluster, shown in grey in Figure 2A. The graphs show distribution, boxplot, mean (black dot and line) and individual paired values. We found descriptive higher decoding accuracy in the auditory ROI for the decoding between visual-manual and auditory-vocal tasks (modality compatible) compared to the decoding between visual-vocal and auditory-manual tasks (modality incompatible). All pairwise t-tests are corrected with the Benjamini-Hochberg procedure (Benjamini & Hochber, 1995). **B**: We controlled for task difficulty to ensure that diffiuclty is not the primary source for the classifier. Here we trained the classifier with a cross-classification again on group-based spheric ROIs (orange dot in Figure 2A) to distinguish visual-manual-easy and auditory-vocal-difficult and tested on visual-manual-difficult and auditory-vocal-easy. If differences in difficulty are the primary source of task classification, the crossing would lead to systematic confusion (i.e., below-chance accuracies), which is clearly not the case. We found significant higher decoding accuracies in the auditory ROI for the modality compatible decoding compared to the modality incompatible decoding. **C**: We also controlled for the modality of task instruction, which was either an image or text. We applied the same cross-classification, which again showed that the primary source of the classification is not the type of instruction. Consistent with the other analysis we again found significant higher decoding accuracies in the auditory ROI for the modality decoding compared to the modality incompatible decoding.

**Figure 4.**
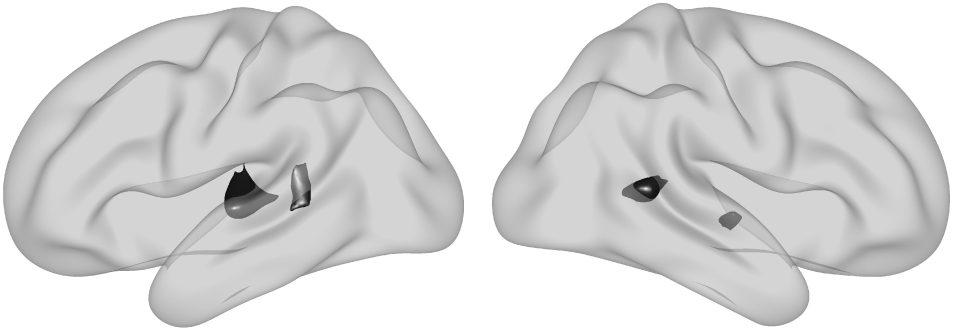
WholeBrain searchlight. *Note:* In addition to all ROI-based analysis, we also implemented a 11mm radius whole brain searchlight to ensure that we captured all relevenat brain regions. The whole brain searchlight was applied separately to each modality mapping, then the difference maps between compatible and incompatible were tested against zero. The resulting cluster remained after FWE correction with *p* = .05, which highly overlaps with the auditory region used as ROI.

In sum, these findings demonstrate that the neural task representations of the two single tasks in the modality incompatible mapping overlap more in the auditory regions than in the modality compatible mapping, supporting the assumed difference in sensory task representation for modality compatible and modality incompatible tasks. Thus and for the first time, we provide evidence that the theoretical overlap of stimulus and action-effect modalities is also represented in neural single-task representations in sensory regions instead of general multitasking-related fronto-parietal regions.

### Practice-related Changes of Neural Task Representations and their Relation to Multitasking Performance

To further substantiate the role of sensory neural overlap for multitasking performance in modality mappings, we examined whether the difference in overlap in the auditory region changes with practice and if this change is related to behavioral change. As the functional organization of the brain is highly variable between individuals, we here used the individual maximum voxel within each group cluster in the localizer task to define individual spheric ROIs for these Pre-Post comparisons (Figure 2B). Performance gains and changes in decoding accuracy were defined as the difference between Pre and Post-timepoint; higher values indicate a performance gain and an increase in decoding accuracy after the practice intervention, respectively.

We did not find a significant effect of time point in the task representation after the practice intervention for any practice group, (main effect timepoint, *F* (1, 60) = 0.01, *p* = .928, 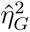 = .000, 90% CI [.000*, .*000], interaction timepoint and intervention group, *F* (2, 60) = 0.61, *p* = .547, 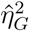 = .005, 90% CI [.000*, .*039]), indicating that on the group level, task representations were relatively stable over time after practice. However, we found a significant positive correlation between the individual changes in decoding accuracy and the performance gains, specifically for the modality incompatible group, *r* = .61, 95% CI [.20*, .*84], *t*(16) = 3.06, *p* = .007, (Figure 2B). A similar correlation was found in the group-specific ROI (*r* = .45, 95% CI [*−.*02*, .*76], *t*(16) = 2.01, *p* = .062) and group-specific cluster (*r* = .55, 95% CI [.11*, .*81], *t*(16) = 2.61, *p* = .019). We additionally applied a multiple regression analysis on the individual ROI to rule out that single-task performance or head movement is responsible for the association (Garner & Dux, 2015). We built a model to explain the performance gain in the modality incompatible mapping for the modality incompatible group, including the single task performance, head movement (as framewise-displacement) and change in decoding accuracy, each for both timepoints and separately for the two intervention groups. The comparison of the beta coefficients of the models revealed that the multitasking improvements in the modality incompatible mapping are predicted to a greater extent for the incompatible intervention group, compared to the compatible intervention group (*β* = 0.42 vs. *β* = -0.23, *z* = 19.50, *p < .*001).

This result confirmed that the significant correlation is not due to differences in single-task performance or head movement. Accordingly, the degree of separation of task representations in auditory regions after practicing a modality incompatible mapping for one session can be considered an important predictor for the elimination of modality-specific dual-task interference within a given session.

## Discussion

While multimodality is a typical characteristic of most everyday multitasking situations, little is known about modality-specific multitasking costs, going beyond attentional or motor limitations. Here, we investigated how modality-based crosstalk between action-effect modality and stimulus modality (Schacherer & Hazeltine, 2020) evolves on a neural level. Specifically, we examined whether modality-specific neural overlap is coded in general multitasking-related brain regions or modality-specific sensory regions. We further elucidated how it affects multitasking performance, and how practice changes those representations. In line with the modality-based crosstalk assumption (Hazeltine et al., 2006; Schacherer & Hazeltine, 2020), we found a significant difference between modality compatible and modality incompatible single-task representations in modality-specific sensory brain regions and not in multitasking-related regions in frontal and parietal cortex. In addition, practice-related improvements in modality incompatible decoding accuracy were associated with performance gains in the modality incompatible dual task. Individuals who succeeded most in reducing modality-specific dual-task costs were those with greatest sensory separation, supporting the assumed relevance of sensory representations for multitasking performance. This effect was only present for the group who practiced the modality incompatible mapping during the intervention, suggesting the build-up of highly specific task representations to deal with potential crosstalk.

For the first time, we provide evidence from neural data for the relation between dual-task crosstalk and sensory modalities, specifically in the auditory cortex. This complements previous findings which revealed multitasking training-related changes of representational overlap in fronto-parietal regions (Garner & Dux, 2015). Garner & Dux exclusively investigated dual-task in relation to single-task performance, thus addressing the representational basis of multitasking costs per se. In our study, in contrast, we directly compared dual tasks with different degrees of modality overlap, addressing specifically the basis of modality-based crosstalk in the context of modality compatibility (Hazeltine et al., 2006; Stelzel et al., 2006). While reducing representational overlap in fronto-parietal cortex may improve the general ability to concurrently process two tasks simultaneously (Garner & Dux, 2015; Garner & Dux, 2023), reducing representational overlap in modality-specific regions may reduce modality-specific sources of multitasking-costs such as modality-based crosstalk.

Our findings are in line with several studies on the role of sensory and motor regions for the representation of stimuli and/or responses and fronto-parietal cortex for representing task rules (see review Woolgar et al. (2016)). Our data support the importance of the fronto-parietal cortex to distinguish between single tasks (significant above chance decoding), but the decoding seems to be independent of the modality mappings. Previous dual-task research on the role of the fronto-parietal cortex was mostly focusing on processes involved in dual-tasking, applying mainly univariate analyses (see review Worringer et al. (2019)). Integrating these findings into representational approaches will be a challenge for future studies in the field. Likewise, asymmetries in modality-specific representations need to be considered in more detail. ln the present study, modality-specific effects were exclusively present in the auditory cortex without comparable effects in visual regions. One potential explanation for the significance of auditory brain regions associated with anticipated action effects of vocal responses is provided by the forward model for self-initiated movements. Following this model, sensory cortices receive a copy of a motor command (i.e., efference copy) while planning a movement and its effects (i.e., action effect) (Holst & Mittelstaedt, 1950; Ody et al., 2023; Wolpert, 1997). Several studies demonstrated high modulation of the auditory cortex as a consequence of speech. More specifically, already the anticipated auditory signal from speech changes activity in the auditory cortex when actually hearing it (Ford et al., 2005; Heinks-Maldonado et al., 2005; Niziolek et al., 2013). In contrast, Straube and colleagues used self-generated button presses and manipulated the multisensory consequences (visual and auditory). They provided evidence that a button press leads to a general preparation to process any following stimulus, irrelevant of the modality. In other words, the expected sensory outcome of a manual button press seems to be broader than the auditory action effect of a vocal response. This is also reflected in everyday experiences where pressing a button can result in a visual effect (i.e., turning the light on and off) or in an acoustic effect (i.e., pressing the doorbell). Whereas speech always results in an auditory effect. It might be that the action effects of a button press are more distributed over the cortex and thus more difficult to decode in sensory regions. On the other hand, the specific action effect of speech can reliably be decoded in the auditory cortex.

Importantly, action effects are a matter of learning. According to the ideomotor theory, the process of action selection is based on the sensory effects of this action, suggesting that there is a bidirectional connection between the action and the action effects (Greenwald, 1970; James, 1890; Prinz, 1997). Several studies provided evidence that the association between action and the action effect is not necessarily hard-wired but can be learned and thus affect task performance (Kühn et al., 2010; Schacherer & Hazeltine, 2021, 2023). For example Kühn and colleagues established images of faces as an artificial action effect of one button press and images of houses for another button press. After this practice phase, the button press was sufficient to activate the neural representations of the previously paired types of images without actually presenting them (Kühn et al., 2010). They concluded that not only action effects guide action selection, but also could an action itself activate a corresponding perceptual representation.

Our study provides additional evidence that the connection between action, action effect and stimulus is relevant for task performance and can be changed, even during a comparatively short practice intervention. Consequently, we assume that in the modality incompatible intervention group, participants have learned to transiently overwrite highly learned modality-specific associations presumably by suppressing the interfering action effect representation and/or by building up a new one. This led to increased performance after the practice intervention, associated with better decodability between single-task representations and decreased performance for the non-practiced modality compatible mapping. Future studies may address the dynamics of suppression and/or building a new association in more detail, and explore further how stable those associations are across time. Taken together, we could provide evidence that not only fronto-parietal regions but also sensory regions hold information about task representations, including action effects, which may be subject to crosstalk in a multisensory multitasking context. These findings reveal for the first time in humans that the neural representation of tasks in a multimodal setting is malleable through multitasking practice at the individual level.

## Methods

This study was pre-registered prior to data analyses (https://osf.io/whpz8). Accordingly, sections in the methods are mostly copied from the preregistration and shortened. We explicitly report any deviations.

### Participants

The total sample of this study consisted of 71 healthy right-handed adults aged 18 to 30 years with German as their first language (or comparable level) and normal or corrected-to-normal vision. Exclusion criteria were any neurological or psychiatric diseases, current medical conditions that could potentially influence brain functions, past or present substance abuse (alcohol and drugs), a self-reported weakness in distinguishing left and right, and common contraindications for MRI scanning. Participants were excluded from the specific analysis if their head movement exceeded the threshold of 25% volumes with framewise displacement > 0.4 mm or if they committed more than 30% errors per run in more than three single task runs (one localizer run), or if the error rate during the practice intervention was higher than 50%. For the dual-task performance, due to the high error rate, we deviated from the pre-registration protocol and limited the 30% criteria to trials in which both stimuli were presented on the same side (averaged for both modality mappings) to ensure that participants were still on task, as incongruence of stimulus information between tasks enhanced task difficulty in addition to modality compatibility (error rate for congruent stimuli: *M* = 17.25, *SD* = 14.90, incongruent stimuli: *M* = 44.92, *SD* = 30.04). In Table 4 we listed the specific exclusion numbers and reasons for each analysis. All three groups were very similar in age and gender distribution (∼ 50% female) (see Table 3). All participants gave their written informed consent before the first session of the study and could choose between 60 € or course credit for reimbursement after completing all sessions. The ethics committee of the Freie Universität approved the study following the Declaration of Helsinki.

**Table 3.** Participants age and sex ratio per intervention group and analysis.

|  | Behavioral analysis |  |  | Decoding analysis Pre timepoint |  |  | Correlation between behavioral and decoding data |  |  |
| --- | --- | --- | --- | --- | --- | --- | --- | --- | --- |
|  | Compatible Intervention | Incompatible Intervention | Passive Control | Compatible Intervention | Incompatible Intervention | Passive Control | Compatible Intervention | Incompatible Intervention | Passive Control |
| Age | 22.33<br>(2.70) | 22.80<br>(2.26) | 22.16<br>(3.47) | 22.17<br>(2.71) | 22.88<br>(2.68) | 22.32<br>(3.27) | 22.24<br>(2.75) | 22.61<br>(2.25) | 22.16<br>(3.47) |
| Female:Male | 7:11 | 10:10 | 9:10 | 11:12 | 13:11 | 12:10 | 7:10 | 8:10 | 9:10 |
| N | 18 | 20 | 19 | 23 | 24 | 22 | 17 | 18 | 19 |

Showing means (standard deviation) for age and number of participants.

**Table 4.**
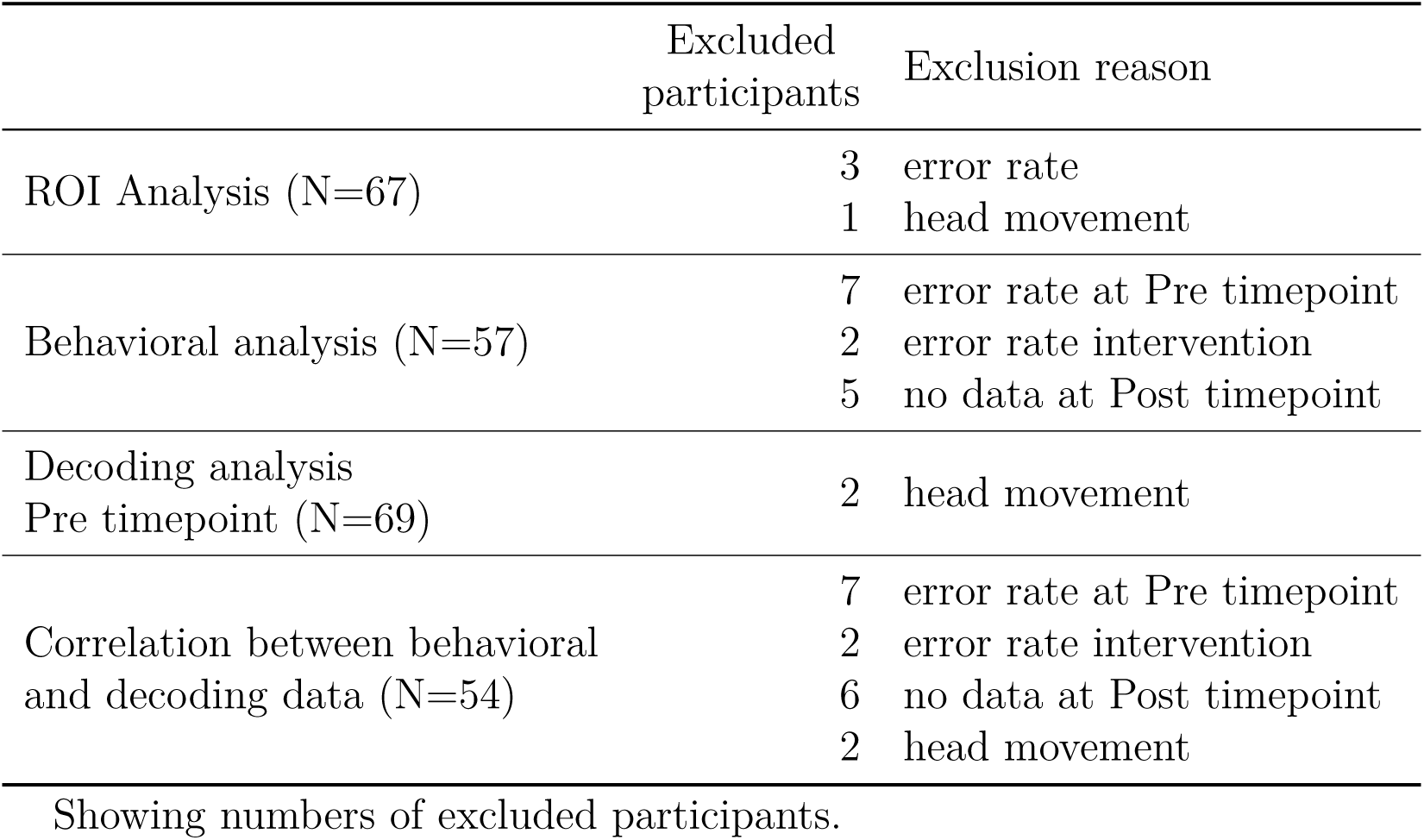
Sample size and exclusion reasons per analysis.

**Table 5.** Center coordinates for spheric ROIs in MNI space.

| ROI Modality | cluster size | x | y | z |
| --- | --- | --- | --- | --- |
| auditory | 1793 | -61 | -31 | 14 |
|  | 3139 | 66 | -19 | 10 |
| visual | 5770 | -27 | -79 | 22 |
|  | 6269 | 24 | -97 | 10 |
| manual | 372 | -53 | -27 | 50 |
|  | 210 | 52 | -23 | 48 |
| vocal | 1995 | -47 | -13 | 38 |
|  | 5512 | 50 | -7 | 38 |
| frontal | 2766 | -29 | 4 | 62 |
|  | 1418 | 26 | 2 | 62 |

MNI-coordinates of peak activation in each cluster. Cluster size in voxel.

### Experimental Overview

Participants completed three sessions; the first session was held online and included behavioral and cognitive measures which will be reported elsewhere. The remaining two sessions (each 2.5-3 h) took place at the Cognitive Center for Neuroscience Berlin (CCNB). During the second session (Session 1 in Figure 1A), participants started outside the scanner with a short familiarization of the tasks (256 trials, 32 per single task, 64 per dual task) before they completed the session in the MRI. Participants returned to the CCNB for the final session (Session 2 in Figure 1A) after a minimum of five and a maximum of nine days at the same time during the day. They repeated the shortened familiarization (128 trials, 16 per single task, 32 per dual task) and continued with the group-specific practice intervention for 80 min before they finished the study with the Post part in the scanner.

### Behavioral Tasks

Participants performed sensorimotor choice reaction tasks, either as single or dual tasks, with varying modality mappings (modality compatible or modality incompatible, compare Figure 1B). In the visual domain, the presented stimuli were a white square (pixel size 56.8 x 56.8) on a black background at six different positions (top, center, bottom), three on the right side of a white fixation cross (pixel size 41.1 x 41.1, thickness 9.9 pixels) and three on the left. In the auditory domain, stimuli were pure tones in three different frequencies (200, 450, and 900 Hz), presented on either the right or the left ear. In the dual-task blocks, stimuli were presented simultaneously. Participants had to respond to the side of the stimuli by either pressing a button with their right or left hand (index finger) and/or by saying the German word for “right” and “left”. The pairing of stimulus and response modality determines the modality mapping. The combination of visual-manual and auditory-vocal is considered modality compatible and the combination of visual-vocal and auditory-manual modality incompatible. Consequently, there was no overlap in either response or stimulus modality within each dual-task condition.

Additionally, we manipulated the task difficulty in the single-task runs by adding visual noise to the stimuli, increasing the distance between the fixation cross and the stimulus, and reducing the contrast between the stimulus and background. For the auditory stimulus, we also added white noise and reduced the volume of the tone compared to the noise. We only included the easy blocks for the behavioral analysis and used the difficulty manipulation as a control analysis for the MVPA. Stimulus material is provided online (https://osf.io/w9hsu/). We randomized per participant the order of the dual-task runs (first modality compatible vs. first modality incompatible mapping) and the block position for the single tasks for each run. Within each block, each stimulus was presented equally often and in random order. To prevent a systematic confound from the visual appearance of the task instruction that was shown at the beginning of each block, we showed task instructions as either a small picture or a text, with different pictures and different fonts for each block to prevent any repetition in visual appearance. Deviating from the Preregistration, we decided against the separate analysis for reaction times and accuracy rates but used an integrated balanced score BIS (Liesefeld & Janczyk, 2019). Reaction times and accuracies are z-standardized and subtracted, resulting in the final BIS parameter. All statistical analysis and plotting was done in R [version 4.2.2; R Core Team (2020)] with RStudio [version 2023.12.1; RStudio Team (2019)].

### Practice Intervention

The practice intervention was completed outside the scanner and consisted only of dual-task trials. For the compatible intervention group, only modality compatible dual-task trials were presented, and for the incompatible intervention group, only modality incompatible dual-task trials. Participants worked on seven runs, each consisting of four blocks with 64 trials per block (a total of 1792 trials). After each run, participants were asked about their subjective feelings in terms of focus, motivation, fatigue, and frustration. The stimulus presentation was the same as for the Pre- and Post measurements. After completing the intervention, participants entered the scanner and completed another run of the intervention tasks (total of 256 trials), inside the scanner without measuring the brain activity. Participants in the passive control group paused for 80 minutes and started immediately with the Post part in the scanner.

### MRI Session

The first fMRI session consisted of a resting state scan and twelve task runs in a block design. Participants started with a 10-minute resting state scan with eyes open in the scanner, followed by two runs of a localizer task, used to define the regions of interest (ROI), each including single and dual tasks in both modality mappings. Each run of the localizer tasks contained six blocks, while each block included 16 trials. In the Pre part the first two runs contained only dual-task trials, and each run was assigned to one modality mapping, consisting of 128 trials. The remaining eight runs contained only single-task trials in both modality mappings, with an easy and a difficult version of the tasks. In each run, every task, modality mapping, and difficulty combination occurred only once, resulting in eight blocks with 16 trials per block. All stimuli were presented for 200 ms, followed by a response interval of 1500 ms and an inter-stimulus interval of 200 ms. Each run concluded with an 8 sec fixation period. The session lasted about 2.5 hours. The Post fMRI session after the practice intervention was the same as the Pre session, starting with the two dual-task runs, followed by eight single-task runs. The session lasted about 3 hours.

### MRI Data Acquisition

Due to a scanner upgrade at the imaging center, the data was acquired with two different scanners. For both scanners, the same head coil and parameters were used. Each participant completed both sessions in the same scanner. The first 25 participants (10x passive intervention, 6x modality compatible intervention, 6x modality incompatible intervention, 3x only Pre-measurement) were measured with Siemens Magnetom TIM TRIO syngo 3T and the remaining participants with Siemens Magnetom 3.0T Prisma both with a 32-channel head coil. At the end of the first session, a high-resolution T1-weighted structural image was measured with 176 interleaved slices, 1 mm isotropic voxels; TE = 2.52 ms, TR = 1900 ms, FoV = 256 x 256 x 176 mm. Functional runs consisted of 139 whole-brain echo-planar images of 37 interleaved slices for the localizer task and the dual-task runs, and 183 whole-brain echo-planar images for each single task run. Each functional run was acquired with 3 mm isotropic voxels, TE = 30 ms, TR = 2000 ms, flip angle =75°, FoV = 192 x 192 x 133 mm. After each dual-task run, a grey-field mapping was measured (3 mm isotropic voxel, TE1 = 4.92 ms and TE2 = 7.38 ms; TR = 400 ms; FoV = 192 x 192 x 133 mm, flip angle = 60°). Participants received auditory stimuli via MRI-compatible headphones (Sensi-Metrics S14, SensiMetrics, USA). Visual stimuli were projected on a screen at the end of the bore, which participants could view through a mirror attached to the head coil. Their vocal responses were recorded via an MRI-compatible microphone (OptimicTM MEG, Optoacoustics, Israel) and the manual responses via MRI-compatible 4-button bimanual boxes (HHSC-2x2, Current Designs, USA).

### MRI Univariate Data Analysis and ROI Definition

Data was converted into BIDS format using dmc2bids (Version 2.1.6; Boré et al. (2023)) and preprocessed using fMRIprep [Version 21.0.2; Esteban et al. (2019), including 3D motion correction and slice-time correction. BIDS transformed raw data was uploaded to OpenNeuro (https://doi.org/10.18112/openneuro.ds005038.v1.0.1). All functional data were aligned to a generated reference image, co-registrated, and transformed to standard space. Anatomical T1-weighted data was resembled into standard MNI space. For more details, please see the generated output script provided by fMRIprep in the preregistration. BOLD runs of the localizer task were smoothed with SPM12 and 8 mm FWHM Gaussian. We used SPM12 to conduct the first-level analysis on all normalized BOLD runs using a block design and a general linear model separately for the localizer runs and the single task runs, the latter also separate for each timepoint (Pre and Post-timepoint). In the localizer model, we included six motion parameters (3x rotation, 3x translation) and the combined measure for head movement framewise displacement as regressors of no interest. For each participant, statistical parametrical maps with contrasts between the stimulus modalities (visual vs. auditory), response modalities (vocal vs. manual), and single vs. dual-task were generated. For the group analysis, the individual maps were averaged and voxelwise tested with a one-sample t-test between the defined contrasts. A cluster-wise FWE-corrected significant threshold (*p* = .05) on the voxel level was used. Note that we restricted the cluster selection to the frontal lobe for the single vs. dual-task contrast. Beta-images used to define clusters were uploaded to Neurovault (https://identifiers.org/neurovault.collection:16842). The resulting highest activation clusters for each contrast (visual, auditory, manual, vocal, and frontal) were used to define the ROI. The clusters served as boundaries to determine the group-based activation peak. This voxel was defined as the center for a 10 mm sphere. A group-based sphere for each contrast was defined as well. We included post hoc additionally an individual differences approach and also identified individual peaks within the group clusters. In preparation for the MVPA analysis, for the single-task model, we included only regressors of each single-task combination (visual-manual, visual-vocal, auditory-manual, auditory-vocal, each in both difficulty levels) but without motion as a regressor.

### Multivoxel Pattern Analysis

After performing the first-level analysis on the single-task runs, we submitted the resulting subject-specific beta images to the Decoding Toolbox (Hebart et al., 2014) to create individual decoding maps for each modality mapping. We used the default methods in the Decoding Toolbox, such as the support vector machines (SVMs) as a decoding method, a leave-one-run-out-cross-classification, and an ROI analysis. This resulted in one decoding accuracy value for each ROI (auditory, visual, vocal, manual, frontal) and each modality mapping (modality compatible and modality incompatible) with a chance level of 50%. We compared our results with the spheric ROI defined by the group activation and with the whole activation cluster as ROI to ensure that our results are not dependent on the small spheres (compare Figure 3. We also included a whole brain searchlight (radius 11 mm) to rule out the option that other brain regions not identified in the univariate analysis contained information about the modality mapping. This was not the case. Only auditory regions were significantly different for the two modality mappings (Figure 3). To further ensure that our results were not influenced by differences in task difficulty and type of instruction (instruction as text or image), we employed a cross-classification on the different difficulty levels and types of instructions, respectively. Specifically, we trained the classifiers to decode between visual-manual-easy vs. auditory-vocal-difficult single tasks and tested on visual-manual-difficult vs. auditory-vocal-easy (respectively for the modality incompatible mapping). This procedure eliminates the influence of task difficulty on the decoding accuracy between the two tasks. In both analyses, the difference between the modality mappings in the auditory regions was still significant (pairwise t-test, corrected with the Benjamini-Hochberg procedure (Benjamini & Hochberg, 1995)). Those results ruled out the explanation that the classifier only differentiated between difficulty or instruction type.

## Acknowledgements

We thank Elisa Arnold, Friederike Glueck, Gregory Gutmann, Lea Lowak, Max Nowaczyk and Oliver Stegmann for assisting in data collection and preprocessing of the vocal data. Neuroimaging was performed at the Cognitive Center for Neuroscience Berlin and was technically supported by Christian Kainz and Till Nierhaus. This work was financially supported by the German Research Foundation, Priority Program SPP 1772 [grant numbers: STE 2226/4-2; GR 3997/4-2; HE 7464/1-2; RA 1047/4-2].

## Notes

### Competing Interest Statement

The authors have declared no competing interest.

